# Development and validation of inducible protein degradation and quantitative phosphoproteomics to identify kinase-substrate relationships

**DOI:** 10.1101/2021.12.08.471812

**Authors:** Rufus Hards, Charles L. Howarth, Kwame Wiredu, Ian LaCroix, Juan Mercado del Valle, Mark Adamo, Arminja N. Kettenbach, Andrew J. Holland, Scott A. Gerber

## Abstract

Phosphorylation signaling is an essential post-translational regulatory mechanism that governs almost all eukaryotic biological processes and is controlled by an interplay between protein kinases and phosphatases. Knowledge of direct substrates of kinases provides evidence of mechanisms that relate activity to biological function. Linking kinases to their protein substrates can be achieved by inhibiting or reducing kinase activity and quantitative comparisons of phosphoproteomes in the presence and absence of kinase activity. Unfortunately, most of the human kinases lack chemical inhibitors with selectivity required to unambiguously assign protein substrates to their respective kinases. Here, we develop and validate a chemical proteomics strategy for linking kinase activities to protein substrates via targeted protein degradation and quantitative phosphoproteomics and apply it to the well-studied, essential mitotic regulator polo-like kinase 1 (Plk1). We leveraged the Tir1/auxin system to engineer HeLa cells with endogenously homozygous auxin-inducible degron (AID)-Plk1). We used HeLa cells and determined the impact of AID-tagging on Plk1 activity, localization, protein interactors, and substrate motifs. Using quantitative proteomics, we show that of over 8,000 proteins quantified, auxin addition is highly selective for degrading AID-Plk1 in mitotic cells. Comparison of phosphoproteome changes in response to chemical Plk1 inhibition to auxin-induced degradation revealed a striking degree of correlation. Finally, we explored basal protein turnover as a potential basis for clonal differences in auxin-induced degradation rates for AID-Plk1 cells. Taken together, our work provides a roadmap for the application of AID technology as a general strategy for the kinome-wide discovery of kinase-substrate relationships.

## Introduction

Phosphorylation signaling is an essential post-translational regulatory mechanism that governs almost all eukaryotic biological processes. In humans, phosphorylation signaling is controlled by the dynamic interplay between protein kinases and their opposing phosphatases on substrate proteins and is frequently deregulated in diseases such as cancer, diabetes and neurodegeneration. Targeting phosphorylation signaling has emerged as a powerful therapeutic strategy in many of these diseases [1-3]. However, challenges in target validation and inhibitor selectivity have limited the clinical use of inhibitors to a small subset of kinases [4]. Kinome-wide RNAi screens in cancer cell lines found that proliferation and survival of these cells depend on a diverse array of kinases, many of which are un- or under-studied [5, 6]. Surprisingly, the degree of overlap for these “essential” kinases is low between cancer cell lines of different tissue origin and/or mutational profiles [4]. While powerful, approaches that rely on systematic genome sequencing, gene expression or RNAi do not elaborate the specific signaling pathways or direct molecular targets.

To achieve a comprehensive understanding of cellular signaling in basic biology and translational medicine, researchers have sought to connect the activities of kinases to the specific phosphorylation sites on their substrate proteins. Approaches to accomplish this goal include the use of kinase inhibitors [7-11], ATP analog-sensitive mutations [12-15], and mRNA silencing [16] to abrogate kinase signaling, followed by quantitative phosphoproteomics analysis to identify phosphopeptides of reduced abundance compared to untreated control phosphoproteomes (**Figure 1A**). However, the availability and selectivity of kinase inhibitors is limited: for example, for the closely related family of Cyclin-dependent kinases (Cdks), the early inhibitor Flavopiridol inhibits Cdk1, 2, 4, 6, 7, and 9 [17-19]. Extensive drug discovery efforts were required to generate more selective inhibitors of Cdk1 [20] and Cdk4/6 [21]. While conceptually innovative, mutations that render a target kinase sensitive to ATP analogs frequently affects their activity, stability, and solubility, and often, suppressor mutations do not restore activity [13]. Finally, while broadly applicable, mRNA silencing requires prolonged periods of treatment, resulting in secondary changes in phosphorylation signaling downstream of the kinase of interest.

**Figure 1.**
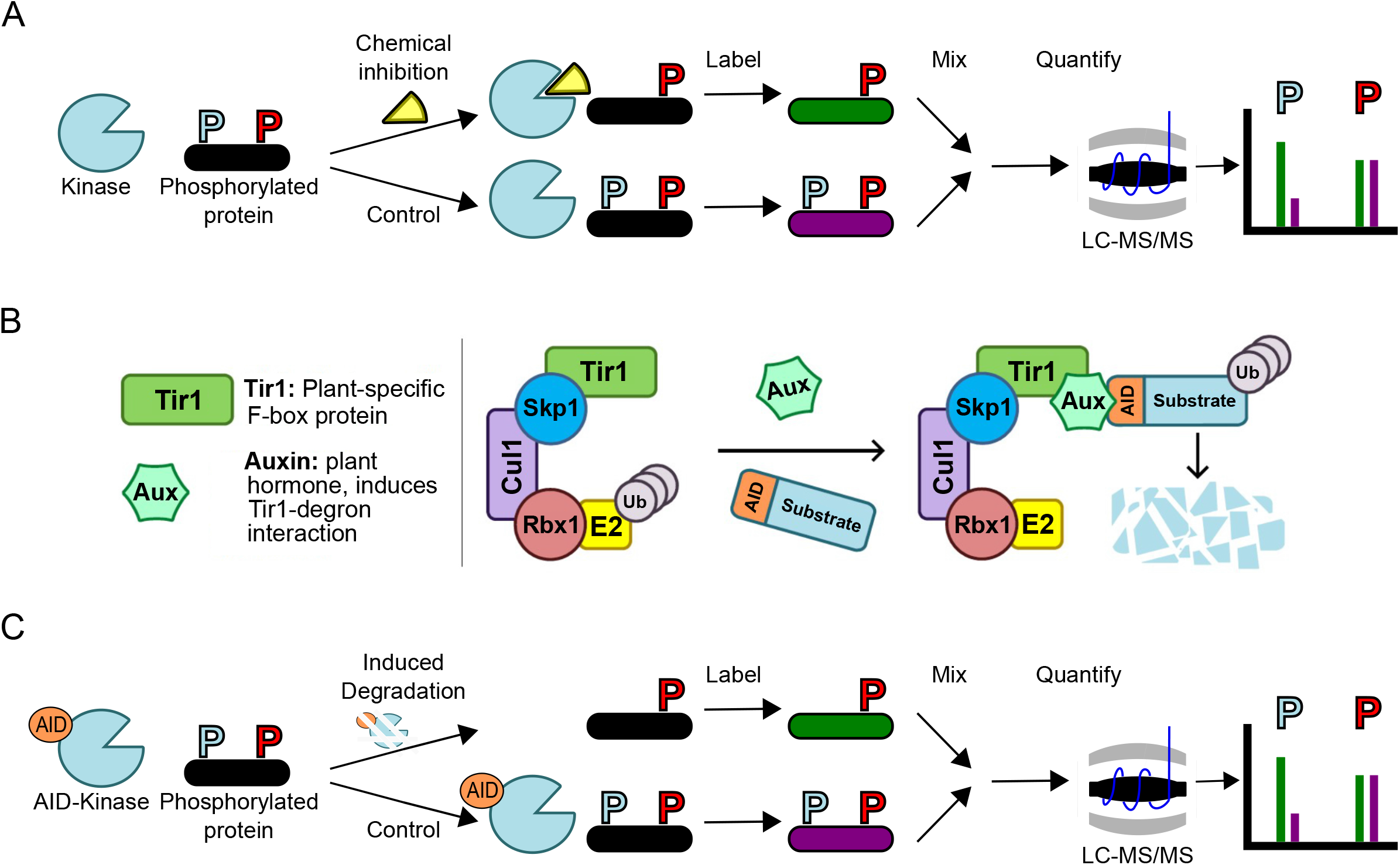
Phosphoproteomic approaches to kinase-substrate discovery. (A) Chemical inhibition of a kinase of interest can uncover substrate phosphorylation sites through comparative quantitative phosphoproteomics. Two cell populations are differentially treated with selective inhibitor or vehicle, and the two conditions are labeled before mixing and analysis for phosphopeptide abundance by mass spectrometry. Candidate substrate sites (blue) should exhibit a decrease in abundance due to inhibition, while non-substrate sites (red) should remain unaffected. (B) Schematic depiction of auxin-inducible degron (AID) system. Plant Tir1 F-box protein is engineered into mammalian cells, where it forms an active E3 ubiquitin ligase complex. This complex ubiquitylates and induces the selective degradation of an AID tagged protein upon addition of the plant hormone auxin. (C) AID-tagged kinases function analogous to but distinct from chemical inhibition, insofar as kinase activity is abrogated by selective degradation upon addition of auxin.

In the present work, we sought to address these issues by combining targeted protein degradation via the Tir1/auxin system [22, 23] with quantitative phosphoproteomics to establish kinase-substrate relationships. In plants, the F-box protein Tir1 recognizes a specific sequence in protein substrates only in the presence of the plant hormone auxin and subsequently targets them for degradation by the proteasome (**Figure 1B**). By fusing the auxin-inducible degron (AID) recognition sequence to a target protein and exogenously expressingTir1, auxin addition to human cells induces the specific and selective degradation of the AID-tagged protein. Thus, depletion of an AID-tagged kinase via auxin addition is analogous to but distinct from chemical inhibition for identifying kinase substrates (**Figure 1C**). Here, we implemented this approach for polo-like kinase 1 (Plk1), an essential mitotic kinase for which we have previously identified candidate substrates via chemical inhibition and quantitative phosphoproteomics [9].

Plk1 activity promotes Cdk1/cyclin B activation and thereby mitotic entry, regulates centrosome separation and maturation as well as spindle assembly, is required for the removal of sister chromatid cohesion and spindle checkpoint signaling, and fine-tunes microtubule-kinetochore attachments and their dynamics [24-27]. Plk1 also contributes to mitotic exit and cytokinesis by recruiting proteins to the central spindle and the midbody, and by regulating the anaphase promoting complex/cyclosome (APC/C) [28, 25, 26]. We generated endogenously tagged AID-Plk1 cell lines and introduced Tir1 into them. We compared the quantitative responses of phosphorylation sites of these cells upon auxin-induced targeted protein degradation to chemical inhibition using the selective small molecule Plk inhibitor BI2536 [29]. We found that the ability of the Tir1/AID-Plk1 system to replicate chemical inhibition depends strongly on the rate and extent of AID-Plk1 turnover, features that appear unique to individual AID-Plk1 knock-in clones. To determine if AID-Plk1 clones that degrade faster in the presence of auxin also exhibit inherently faster proteome-wide protein turnover rates, we performed additional quantitative proteomics time course experiments in the presence of cycloheximide for fast and slow degrading AID-Plk1 cell lines and report on our observations.

## Methods

### Cell culture

HeLa and HEK293T cells were grown in Dulbecco’s Modified Eagle’s Medium (DMEM; Gibco) supplemented with 10% Fetal Bovine Serum (FBS; Hyclone) and 1% penicillin-streptomycin (100 U/mL and 100 μg/mL; Corning). Cells were incubated at 37°C in 5% CO2.

### Cell cycle synchronization

For mitotic arrest, cells were subjected to a double-thymidine block (2 mM; Sigma) for synchronization at the start of S phase in the cell cycle. Cells were incubated for 18 hours in the presence of thymidine, followed by a washout into thymidine free media for 8 hours and an additional incubation in the presence of thymidine for 12 hours. Cells were released from the second thymidine block for 6 hours prior to the addition of Taxol (100nM) for 12 hours.

### Plk1 interactomes

2xHA-Plk1 and 2xHA-sAID-Plk1 were cloned into pcDNA3.1 and transfected into HEK293T cells using JetPrime (Polyplus) transfection reagent per manufacturer’s protocols. After 24 hours, the transfection mixture was replaced with fresh media, and cells were grown for an additional 24 hours prior to replating into G418-(400μg/mL) containing media and selection for 2 weeks to generate stable cell lines.

2xHA-Plk1 and 2xHA-sAID-Plk1 HEK293T cells were synchronized in mitosis as described above in triplicate 10cm dishes. Mitotic cells were collected by shake-off, washed with PBS and lysed in ice-cold 1.2mL immunoprecipitation buffer (20mM Tris pH 7.5, 0.5mM DTT, 150mM NaCl, 0.5% Triton X-100, 2mM sodium fluoride, 2mM sodium molybdate, 5mM β-glycero phosphate, and EDTA-free Mini-Complete protease inhibitors (Roche)) by sonication using a Branson sonicator with a micro-tip at 14% power for 8s three separate times, cooling on ice for 2 minutes in between each sonication. Lysates were clarified by centrifugation (21,000 x g) for 10 minutes at 4°C, and the clarified lysates were treated with anti-HA antibodies coupled to protein-G Sepharose beads (Pierce) for 3 hours with rotation at 4°C. Immunoprecipitates were washed 4x with ice-cold lysis buffer, then twice with lysis buffer without DTT. Proteins were eluted by addition of 100μL 1% SDS/100mM Tris pH 8 solution at 50°C for 5 minutes in a thermomixer at 1000rpm. Eluted proteins were isolated by SP3 isolation as described previously [30] and digested to peptides in 15 mM ammonium bicarbonate buffer. Peptide digests were desalted on an OASIS μHLB plate (Waters) and analyzed by LC-MS/MS as described below.

### Immunofluorescence

Wild-type or 3xFlag-mAID-Plk1 expressing HeLa cells were fixed with 3% formaldehyde, permeabilized with PBS containing 0.1% Triton-X100 (PBST), and incubated with anti-alpha tubulin (CellSignaling) and either anti-Plk1 (Sigma) or anti-Flag (Sigma) primary and appropriate secondary antibodies (Molecular Probes, Invitrogen) at room temperature. DNA was labeled with Hoechst33342 (Sigma). Images were collected using a 63X, 1.4NA objective on an Axioplan2 Zeiss microscope using OpenLab software.

### CRISPR-Cas9 homologous recombination clone generation

For AID-Plk1 clones, homologous recombination constructs were synthesized in parts as gBlocks (IDT) and assembled in a pBluescript-SK-II+ cloning vector. Homologous recombination (HR) arms were targeted around the start codon of Plk1 based on the UCSC genome browser genomic sequence. Left HR arm was 841 bp and the right HR arm was 740bp. HR insert consisted of one of three resistance cassettes (blasticidin, hygromycin, and Zeocin) followed by a P2A ribosomal skip, 3xFLAG tag, and mAID tag. Three gRNA vectors were constructed in the dual U6-gRNA/CMV-Cas9 expression vector pX330 (Addgene) based upon predicted high scoring guides from CRISPOR.tefor.net, and HR targeting templates were synthesized resistant to all gRNAs used with mutations in the gRNA PAM sequence to prevent gRNA recognition. Cells were plated to be ∼60% confluency in 6-well tissue culture dishes at time of transfection. 1.2ug of targeting vector per well was digested with restriction endonucleases to excise the insert for 1 hour at 37C and then purified with EZNA PCR purification columns. Linearized targeting template was combined with 0.4ug of pX330 gRNA vector and transfected into cells with JetPrime (Polyplus) per manufacturer’s protocols. After 24 hours, transfection mixture was replaced with fresh media and cells were allowed to recover for an additional 24 hours prior to starting selection in blasticidin (10μg/mL), hygromycin (100μg/mL), or Zeocin (100μg/mL). Selection agents and media were changed every 2 days until cell death halted and colonies formed. Individual colonies were harvested with dilute trypsin digestion in PBS and manual picking into 96-well plates using a screening microscope at 4x magnification and a pipette.

Tir1 was introduced into AID-Plk1 clones by either retroviral introduction or AAVS1 CRISPR-Cas9 introduction as previously described [31] and selected with the appropriate antibiotic (neomycin/G418 or puromycin, respectively) to generate individual colonies, which were isolated and harvested as above.

### Screening for AID-Plk1 clones

For genomic screening of inserts, 80% of a confluent 96-well was harvested and lysed in QuickExtract (Lucigen). In short, cells pellets were resuspended in 40ul of QuickExtract reagent, heated at 60C for 7 mins, vortexed well, heated at 90C for 2min, vortexed again, and then stored at 4C. PCRs with primers targeting genomic loci exterior to the homologous recombination arms and primers internal to each resistance cassette were used to confirm site specific insertion of resistance cassettes and the loss of wild-type amplicon.

AID-Plk1 screening primers:

Plk1 left arm forward primer 5’- GCATGCTTGTAATTCCAGCTGCTAGG -3’

Plk1 right arm reverse primer 5’- TGACCCAGCTCCACCCCAGAAG -3’

Blasticidin resistance reverse primer 5’- GGTGGATTCTTCTTGAGACAAAGGC -3’

Hygromycin resistance reverse primer 5’- CGGTGTCGTCCATCACAGTTTGCC -3’

Zeocin resistance reverse primer 5’- GAACGGCACTGGTCAACTTGG -3’

Clones that were positive for tag insertion and loss of wild-type PCR amplicon were grown to the 24-well stage where they were screened for AID-Plk1 homozygosity and tag insertion by western blot with Plk1 and Flag antibodies, respectively. Tir1 clones created from homozygous AID clones were screened for Tir1 insertion by western blot with anti-Myc and Tir1b antibodies. Tir1-positive, AID-Plk1 homozygous clones were finally screened for their capacity to degrade AID-Plk1 by western blot after 2 hour auxin (3-indoleacetic acid, IAA or 1-naphthaleneacetic acid, NAA) treatment with anti-Plk1 and Flag antibodies.

### Flag purification of proteins

Various AID length Plk1, analog-sensitive(as) Plk1, COIL, DEK, NCK1, and FAM54B were cloned into the p3XFLAG-CMV-10 vector. Constructs were transformed into the DH5α strain of *E. coli* for plasmid purification. Plasmids were transiently transfected into 60-80% confluent plates of HEK293 cells using polyethylenimine (PEI). After 24 hours cells were dislodged from culture plates with 0.25% trypsin (Corning), centrifuged and washed with PBS. Cells were resuspended in lysis buffer containing Tris pH 7.5 (50 mM), 1% Triton X-100, sodium fluoride (2 mM), sodium molybdate (2 mM), β-glycero phosphate (5 mM), NaCl (400 mM) and EDTA-free Mini-Complete protease inhibitors. Lysate was sonicated using a Branson sonicator equipped with a micro-tip three times at 15% power for 15 seconds each on ice. Lysate was pre-cleared by centrifugation at 21100 x g for 15 minutes at 4°C. Anti-FLAG affinity gel was washed three times with lysis buffer, added to cleared lysates and incubated for 4 hours at 4°C with rotation. Protein bound resin was washed three times with lysis buffer. Protein was eluted with FLAG peptide (166 μg/mL) in Tris-buffered saline. Eluted material was dialyzed for 15 hours with HEPES pH 7.5 (10 mM), NaCl (125 mM), EDTA (100 μM) and DTT (200 μM). Dialyzed material was aliquoted for single use, snap frozen in liquid nitrogen and stored at -80°C.

### Proteomics and phosphoproteomics TMT experiments

AID-Plk1 Tir1 Hela clones B12-11 and 23r3 were cultured to nine 10cm tissue culture dishes each and arrested at mitosis as described above. Triplicates of dishes were treated with 1μM BI2536 for 1 hour, 1mM 1-napthaleneacetic acid (NAA) for 2 hours, or control treated with DMSO for 2 hours. After treatment, cells were collected and washed with PBS prior to snap freezing in liquid nitrogen. Frozen cell pellets were partially thawed on ice and resuspended in lysis buffer containing urea (8M; AMRESCO), Tris pH 8.1 (50 mM; Alfa Aesar), β-glycerophosphate (2 mM; Sigma), sodium fluoride (2 mM; Fluka), sodium molybdate (2 mM; Sigma), sodium orthovanadate (2 mM; Sigma) and EDTA-free Mini-Complete protease inhibitors (1 tablet per 10mL; Roche). Lysates were sonicated using a Branson sonicator equipped with a micro-tip three times at 15% power for 15 seconds each on ice. Protein concentration of the lysates were determined by BCA protein assay (Thermo). Proteins were reduced with dithiothreitol (DTT, 5 mM; Sigma) at 55°C for 30 minutes, cooled to room temperature and alkylated with iodoacetamide (15 mM; Sigma) at room temperature for 1 hour in the dark. The alkylation reaction was quenched by the addition of another 5 mM DTT. The lysates were diluted 6-fold in 25 mM Tris pH 8.1, and 1 mg sequencing grade trypsin (Promega) per 100 mg total protein was added to each lysate and incubated for 16 hours at 37°C. The digests were acidified to pH ≤ 3 by addition of trifluoroacetic acid (TFA; Honeywell) and any resulting precipitates were removed by centrifugation at 7100 RCF for 15 minutes. The acidified lysates were desalted using a 60mg Oasis desalting plate (Waters). A fraction of each clone 23r3 sample elution was taken corresponding to 40ug of total protein based on the BCA quantification and aliquoted separately for total protein abundance proteomics. The eluates were vacuum centrifuged for 30 minutes at 37°C to evaporate organic solvent. Finally, the samples were snap frozen in liquid nitrogen and lyophilized for 16 hours. The dried clone B12-11 samples and the bulk clone 23r3 samples not saved for total protein screening were enriched for phosphopeptides by following the High-Select Fe-NTA phosphopeptide enrichment kit protocol. Phosphopeptide eluents were vacuum centrifuged to dryness before desalting on a 10mg Oasis desalting plate. Phosphopeptide eluents were split into halves and vacuum centrifuged to dryness.

Phosphopeptide samples from clone B12-11 and peptide and phosphopeptide samples from clone 23r3 were labeled with TMT reporter by resuspending the peptides in 166mM EPPS pH 8.5. 70 ug of the appropriate TMT reagent in acetonitrile (ACN) was added to each sample before they were vortexed and incubated at room temperature for 1hour. TMT label reagents were assigned to samples as follows. B12-11 phosphopeptide: Control 1(Ctrl) – 127N, Ctrl 2 – 127C, Ctrl 3 – 128N, BI2536 treated (BI) 1 – 128C, BI 2 – 129N, BI 3 – 129C, Auxin treated (NAA) 1 – 130N, NAA 2 – 130C, NAA 3 – 131N. 23r3 peptide and phosphopeptide: Ctrl 1 – 127N, Ctrl 2 – 127C, Ctrl 3 – 128N, BI 1 – 128C, BI 2 – 129N, BI 3 – 129C, NAA 1 – 130N, NAA 2 – 131N, NAA 3 – 131C. 5% of each labeling reaction was taken, combined with 5% methanol 0.1% TFA, desalting by STAGE tip, and analyzed by LC-MS/MS by including the TMT label as a dynamic modification in the search to confirm TMT labeling efficiency > 95%. Each individual remaining reaction was frozen at -80C until labeling efficiency was determined. If a reaction showed <95% labeling efficiency, the parent reaction was thawed, an additional 20ug of TMT reagent was added, and the labeling reaction was repeated. Once all channels showed >95% labeling efficiency, the reactions were quenched with hydroxylamine for 10 minutes at room temperature. Samples from each comparison were mixed to form the appropriate multiplex and then acidified with TFA to pH < 3 before desalting on an OASIS 10mg desalting plate (Waters). Eluents were dried by vacuum centrifugation before off-line fractionation by HPLC on a pentafluorophenyl (PFP) column as previously described [32]. The 48 fractions from the HPLC fractionation were concatenated into 24 fractions for the phosphopeptide samples, and 16 fractions for the peptide sample. The B12-11 fractions were analyzed by LC-MS/MS as described above on the Orbitrap Fusion Tribrid instrument, while the 23r3 fractions were analyzed on the Orbitrap Fusion Lumos Tribrid instrument.

### LC-MS/MS analyses

LC-MS/MS analysis was performed on either an Orbitrap Fusion Tribrid mass spectrometer or an Orbitrap Fusion Lumos Tribrid mass spectrometer (ThermoFisher Scientific, San Jose, CA) equipped with an EASY-nLC 1000 ultra-high pressure liquid chromatograph (ThermoFisher Scientific, Waltham, MA). Peptides and phosphopeptides were dissolved in loading buffer (5% methanol (Fisher) / 1.5 % formic acid) and injected directly onto an in-house pulled, polymer coated, fritless, fused silica analytical resolving column (35 cm length, 100μm inner diameter; PolyMicro) packed with ReproSil, C_18_ AQ 1.9 μm 120 Å pore stationary phase particles (Dr. Maisch). Phosphopeptides in 3 μl loading buffer were loaded at 450 bar by chasing on to the column with 8μl loading buffer. Samples were separated with a 120-minute gradient of 4 to 33% LC-MS buffer B (LC-MS buffer A: 0.125% formic acid, 3% ACN; LC-MS buffer B: 0.125% formic acid, 95% ACN) at a flow rate of 330 nl/minute. The instruments were operated with an Orbitrap MS^1^ scan at 120K resolution and an AGC target value of 500K. The maximum injection time was 100 milliseconds, the scan range was 350 to 1500 m/z and the dynamic exclusion window was 15 seconds (+/- 15 ppm from precursor ion m/z). Precursor ions were selected for MS^2^ using quadrupole isolation (0.7 m/z isolation width) in a “top speed” (2 second duty cycle), data-dependent manner. MS^2^ scans were generated through collision-induced dissociation (CID) fragmentation (35% CID energy) and either linear ion trap analysis (Rapid setting) for peptides or Orbitrap analysis at 30K resolution for phosphopeptides. Ion charge states of +2 through +4 were selected for HCD MS^2^. The MS^2^ scan maximum injection time was 60 milliseconds and AGC target value was 60K. For TMT runs, top 8 MS2 peaks were dynamically isolated and further fragmented by higher-collision energy (HCD) at 55% via SPS-MS^3^ for quantification of liberated reporter ions (110 – 500 m/z).

### Peptide spectral matching and bioinformatics

Raw data were searched using COMET against a target-decoy version of the human (*Homo sapiens*) proteome sequence database (UniProt; downloaded 2018; 20,241 total proteins) with a precursor mass tolerance of +/- 1.00 Da [33] and requiring fully tryptic peptides with up to 3 missed cleavages, carbamidomethyl cysteine as a fixed modification and oxidized methionine as a variable modification. For phosphopeptide data, searches were expanded to include the dynamic addition of phosphate to serine, threonine, and tyrosine residues. For TMT-labeled samples, the mass of the TMT reagent (229.162932 Da) was added as a static modification to all peptide N-termini and lysine residues. Phosphorylation of serine, threonine and tyrosine were searched with up to 3 variable modifications per peptide, and were localized using the phosphoRS algorithm [34]. The resulting peptide spectral matches were filtered to ≤1% false discovery rate (FDR) by defining thresholds of decoy hit frequencies at particular mass measurement accuracy (measured in parts per million from theoretical), XCorr and delta-XCorr (dCn) values. Statistical analyses and processing of peptide extracted ion chromatograms (XIC) were performed using the R statistical programming language (http://www.R-project.org).

### In vitro kinase reactions

*In vitro* peptide library kinase assays were carried out in HEPES pH 7.5 (10 mM), MgCl_2_ (20 mM), β-glycerophosphate (5 mM), DTT (0.5 mM) and ATP (1 mM). 50 μg dephosphorylated peptide libraries, prepared as done previously [35], were resuspended in assay buffer and incubated with 0.5μg 3xFlag-mAID-Plk1 or 3xFlag-Plk1 at 30°C overnight. Reactions were vacuum centrifuged to dryness, resuspended in 0.1% TFA and desalted using a C_18_ desalting plate. Phosphopeptide enrichment was performed as indicated above. Eluted phosphopeptides were desalted using the stop-and-go-extraction (STAGE-tip) method [36]. Desalted samples were vacuum centrifuged to dryness and stored at -80°C prior to analysis by LC-MS/MS.

Kinase activity was measured using 250ng each 3xFlag-Plk1, 3xFlag-fAID-Plk1, sAID-Plk1, mAID-Plk1, and as-Plk1 in individual reactions with 50μg aliquots of dephosphorylated peptide libraries and ATP as described above, except reactions were carried out at 30°C for 2 hours, followed by analysis of ATP consumption by ADP-Glo assay per manufacturer’s protocol. Chemiluminesence was measured on a SpectraMAX plate luminometer.

Plk1 substrate phosphorylation reactions were performed with 3xFlag-COIL, 3xFlag-DEK, 3xFlag-NCK1, and 3xFlagFAM54B purified as detailed above. Purified, active 10xHis-Plk1 was received as a gift from the Kettenbach lab as previously described [35]. Four assay conditions were used per ∼1ug candidate substrate: -ATP condition (30mM Hepes, 35mM MgCl_2_), +ATP (30mM Hepes, 35mM MgCl2, 500μM ATP disodium salt), +ATP +Plk1 (30mM Hepes, 35mM MgCl_2_, 500μM ATP disodium salt, 100 ng Plk1), and +ATP +Plk1 +BI2536 (30mM Hepes, 35mM MgCl_2_, 500μM ATP disodium salt, 100 ng Plk1, 1μM BI2536). Reactions were incubated at 37°C for 2 hours. Samples were prepared for LC-MS/MS analysis by the single pot stationary phase sample preparation (SP3) protocol as described previously [30]. The resulting peptides were desalted by STAGE-tip and analyzed by LC-MS/MS, and phosphopeptide abundances were measured as area under the curve of extracted ion chromatograms (XIC) for each identified phosphopeptide.

### Cycloheximide chase proteomics by TMT

Clone B12-11 and clone 23r3 cells were grown in triplicate repeats of nine 6-well culture dishes each. Each set of nine was treated in triplicate with 50 μg/ml cycloheximide (CHX) for 4 hours, 20 hours, or control-treated with DMSO. Cells were harvested by trypsin dissociation, washed with PBS, and then snap frozen in liquid nitrogen. Each cell pellet was thawed on ice, lysed in 1 ml of 50mM Tris pH 8.1 40mM NaCl 1% SDS by sonication, analyzed for protein content by BCA assay, and then reduced and alkylated as noted above. A volume of lysate equivalent to 15ug of protein from the control treated cells from the BCA was taken from each sample for SP3 purification, digestion and TMT labeling. Protein samples were processed by SP3 as above, but with the following alterations: the final two bead washes were done with 100% ACN as opposed to 80% ethanol and the trypsin digestion was performed in 166mM EPPS pH 8.5 as opposed to 50mM ammonium bicarbonate. Peptides in EPPS buffer were removed from the SP3 beads after overnight digest, ACN was added to 10% final concentration, and peptides were directly labeled with 27ug of the appropriate TMT channel reagent as above. Samples were labeled with TMT channels as follows. Replicate one (CHXI) of B12-11 and 23r3 9-plex: Control treated (CHX0) 1 – 127C, CHX0 2 – 128N, CHX0 3 – 128C, 4 hour cycloheximide (CHX4) 1 – 129N, CHX4 2 – 129C, CHX4 – 130N, 20 hour cycloheximide (CHX20) 1 – 130C, CHX20 2 – 131N, CHX20 3 -131C. Replicate two (CHXII) and replicate three (CHXIII) of B12-11 and 23r3 9-plex: CHX0 1 – 126, CHX0 2 – 127C, CHX0 3 – 128N, CHX4 1 – 128C, CHX4 2 – 129N, CHX4 3 – 129C, CHX20 1 – 130N, CHX20 2 – 130C, CHX20 3 -131N. Labeling efficiency checks, quenching, desalting, and off-line fractionation were performed as above. The 48 HPLC fractions were concatenated into 24 fractions for each 9-plex, and the following fractions were analyzed for each on the LC-MS/MS on an Orbitrap Fusion Lumos as above: fractions 2, 5, 7, 8, 11, 12, 14, 15, 17, 20, 21, and 23.

### Cycloheximide chase proteomics data processing

We merged the log-transformed values of all identified proteins from both fast and slow-degrading cell lines for downstream analysis; proteins were required to be present in at least 5 of the 9 control replicates for a given cell line. We performed sample-level outlier analysis using OutlierDM, in R and used Mahalanobis distance to remove samples if their collective protein values deviated from the center of distribution of their corresponding replicates. All timepoints were normalized to controls and expressed as fractional protein abundance remaining. For a given timepoint, we normalized replicates to the same total protein concentration (TPN). Following the per-timepoint normalization among replicates, the effect of experimental batches of samples was assessed and corrected for, using the ComBat algorithm [37].

We compared the rate of protein degradation per hour of experimental time between the 2 cell lines and visualized the distribution of rates at 4-hours, in the last 16 hours of experimental time and for the overall experiment. We also compared the difference in the amount of protein left undegraded, *delta*, between 23R3 and B12-11. This was done by normalizing all control samples to unity as 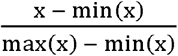 we plotted a histogram of the distribution of *deltas* at both 4-hour and 20-hour experimental timepoints.

## Results

### Shorter auxin-inducible degron (AID) tags do not affect Plk1 localization, protein interactors, kinase activity, or substrate motif preferences

We selected polo-like kinase 1 (Plk1) to develop and validate our approach, a kinase for which we had previously identified candidate substrates *via* chemical inhibition and phosphoproteomics, and performed cell biochemical and biological characterization *in vitro* and in cells [9, 35, 38]. First, we tested the impact of auxin-inducible degron (AID) tagging on Plk1 activity and function by ectopically expressing 3xFlag-tagged Plk1 constructs containing AID tags of various length in HEK293T cells [23, 39]. We also generated an analog-sensitive [40] version of Plk1 (AS-Plk1) by site-directed mutagenesis [13]. We purified 3xFlag-Plk1, 3xFlag-AID-Plk1, and 3xFlag-AS-Plk1 and performed in *vitro* kinase assays on pools of naturally-occurring HeLa peptides as substrates as previously described [35]. We used this to measure the ATP consumption of each kinase version and to determine the linear substrate motif preferences of the different AID-Plk1 variants. While triplicate analysis of full-length AID (fAID, 228 amino acids) tagged Plk1 showed a significant decrease (62%) in ATP turnover relative to 3xFlag-tagged Plk1 alone, shorter AID tags (mAID, 68 amino acids, and sAID, 46 amino acids) showed no significant difference between Flag-tagged only and Flag-AID-tagged kinases (**Figure 2A**). However, AS-Plk1 was significantly reduced in activity by 63% compared to 3xFlag-Plk1. This is consistent with prior reports of AS-Plk1 exhibiting reduced kinase activity [13]. Furthermore, motif enrichment analysis of phosphorylated peptides from *in vitro* kinase reactions with 3xFlag-Plk1 and 3xFlag-sAID-Plk1 revealed indistinguishable substrate preference. Consistent with previous analyses, 3xFlag-Plk1 and 3xFlag-AID-Plk1 preferentially phosphorylated [D/E/N]Xp[S/T][hydrophobic] and p[S/T]F motifs, which are preferred motif sequences for Plk1 phosphorylation [9, 35, 41] (**Figure 2B**). Thus, sAID- and mAID-tags do not negatively impact Plk1 activity or substrate phosphorylation *in vitro*.

**Figure 2.**
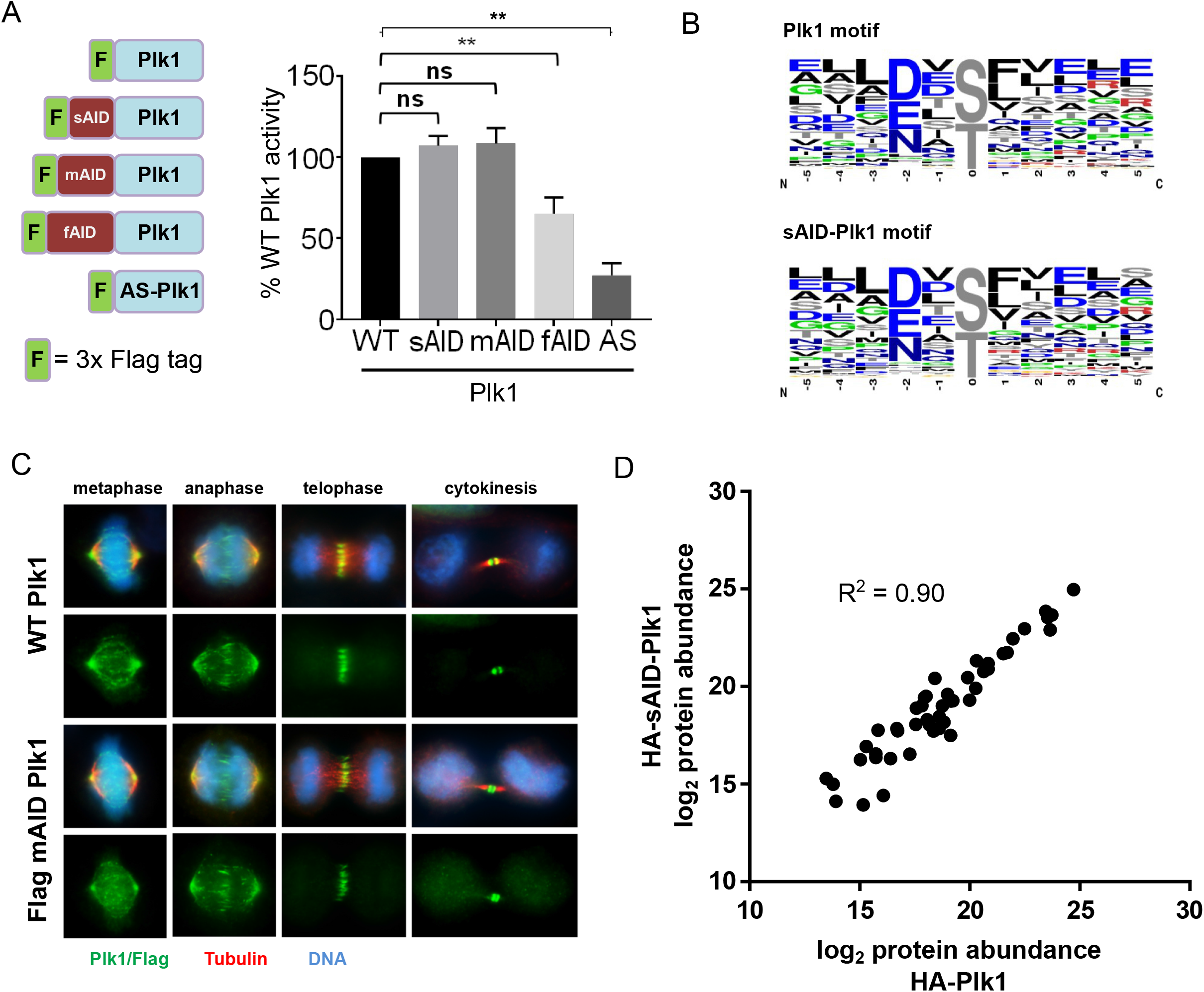
AID tag does not interfere with normal Plk1 function. (A) Kinase activity of Plk1 constructs. 3xFlag (3xF) control, 3xF-sAID (aa 71-114), 3xF-mAID (aa 68-134), 3xF-fAID (aa 1-228), and 3xF-analog-sensitive (AS)-Plk1 were isolated and tested for activity *in vitro*. 3xF-fAID- and 3xF-AS-Plk1 exhibited lower specific activity than wild-type, while both 3xF-sAID- and 3xF-mAID-Plk1 were not significantly different from 3xF-Plk1. (B) Weblogo depictions of kinase motifs generated by *in vitro* kinase activity of 3xF-Plk1 and 3xF-sAID-Plk1 on Hela cell peptide digests. Size of amino acids represents prevalence in phosphorylated peptides over background. (C) Immunofluorescence imaging of Hela cells at different mitotic stages (D) Abundance of known Plk1 polo-box domain (PBD) dependent interactors with HA-Plk1 and HA-sAID-Plk1 from immunoprecipitation and label-free quantitative mass spectrometry.

To determine the effect of AID-tags on Plk1 substrate binding and localization in cells, we generated a HeLa cell line containing 3xFlag-mAID-Plk1. We then subjected it and parental HeLa cells to indirect immunofluorescence microscopy using anti-Flag or anti-Plk1 antibodies, anti-tubulin antibodies to visualize microtubules, and a DNA stain. Plk1 exhibits a distinct, phase-specific localization behavior in mitosis: in prophase cells, Plk1 localizes to spindle poles; in prometaphase and metaphase to spindle poles, spindle microtubules and kinetochores; in anaphase and telophase to the central spindle; and in cytokinesis to the midbody.

Immunofluorescence images of Plk1 demonstrate no differences in localization at any stage of mitosis between 3x-Flag-AID-Plk1 and wild-type Plk1 (**Figure 2C**). Finally, we immunoprecipitated Plk1 from HEK293T cells stably expressing either 2HA-Plk1 or 2HA-sAID-Plk1, arrested in mitosis with Taxol, and analyzed their interactomes by label-free quantitative LC-MS/MS in biological triplicate (**Supplementary Table 1**). Unlike many other kinases, Plk1 stably binds to phosphorylated epitopes on substrates and scaffolding proteins via its conserved polo-box domain [42, 43]. We and others have leveraged this property to identify and validate candidate substrates of Plk1 by protein-protein interaction analysis [44, 9, 38]. Correlation of the average abundances of known Plk1 substrates or phosphorylation-dependent protein interactors between the 2HA-Plk1 and 2HA-sAID-Plk1 interactomes was high (R^2^ = 0.90; **Figure 2D**). Taken together, these data support the notion that shorter AID tags do not alter Plk1localization or protein interactions in cells.

### CRISPR/Cas9-based endogenous tagging to generate homozygous 3xFlag-AID-Plk1 HeLa cells

To generate endogenous AID-tagged Plk1, we designed a homologous recombination (HR) targeting vector with ∼700bp targeting arms that flank the ATG start site of the Plk1 gene locus for inserting a antibiotic resistance gene, a viral ribosomal skip sequence (P2A), and a 3xFlag-sAID sequence in frame with Plk1 (**Figure 3A**). In addition, we identified a CRISPR/Cas9 guide RNA (gRNA) targeting sequence proximal to the start ATG, in the 5’ untranslated region (UTR), and cloned it into a Cas9 and gRNA co-expression plasmid (pX330). We also included a corresponding set of mutations at the gRNA targeting sequence in the HR repair plasmid to prevent cutting of successfully inserted sequences (**Figure 3A**). Initial attempts at endogenous tagging in HeLa cells using this strategy, yielded only single allelic insertions of the AID tag after transfection and selection, even when screening hundreds of individual clones (data not shown). Thus, to generate homozygous clones, we designed three different versions of the HR targeting vector, each containing a unique resistance cassette (blasticidin, hygromycin, or bleomycin). In addition, we identified two additional CRISPR/Cas9 gRNA targeting sequences near the start ATG but at least 30 nucleotides away from any other gRNA sequence and included mutations in all three copies of the targeting plasmids to render each construct transparent to the three separate Cas9-gRNA sites (**Figure 3A**). We then transfected HeLa cells with these constructs in a stepwise fashion followed by antibiotic selection to generate homozygous AID-tagged clones. We observed a progressive increase in the ratio of AID-tagged to wild-type Plk1 by western blot of pooled lysates after each successive round of transfection and selection (**Figure 3B**). After three rounds, individual clones were collected and screened for homozygous insertion of the AID tag by genomic PCR (**Figure 3C**) and confirmed by western blot **(Figure S1)**. We then introduced *os*Tir1 (Tir1) using either CRISPR/Cas9-mediated insertion to the AAVS1 “safe harbor” locus [31] or retroviral insertion randomly into the genome. After selection, we screened single cell clones for Tir1 expression and determined their capacity to degrade 3xFlag-sAID-Plk1 upon addition of auxin for 2 hours (**Figure 3D**). This process generated several clonal HeLa cell lines with homozygous, endogenously tagged 3xFlag-sAID-Plk1 that degraded AID-Plk1 to different extents upon addition of auxin for 2 hours.

**Figure 3.**
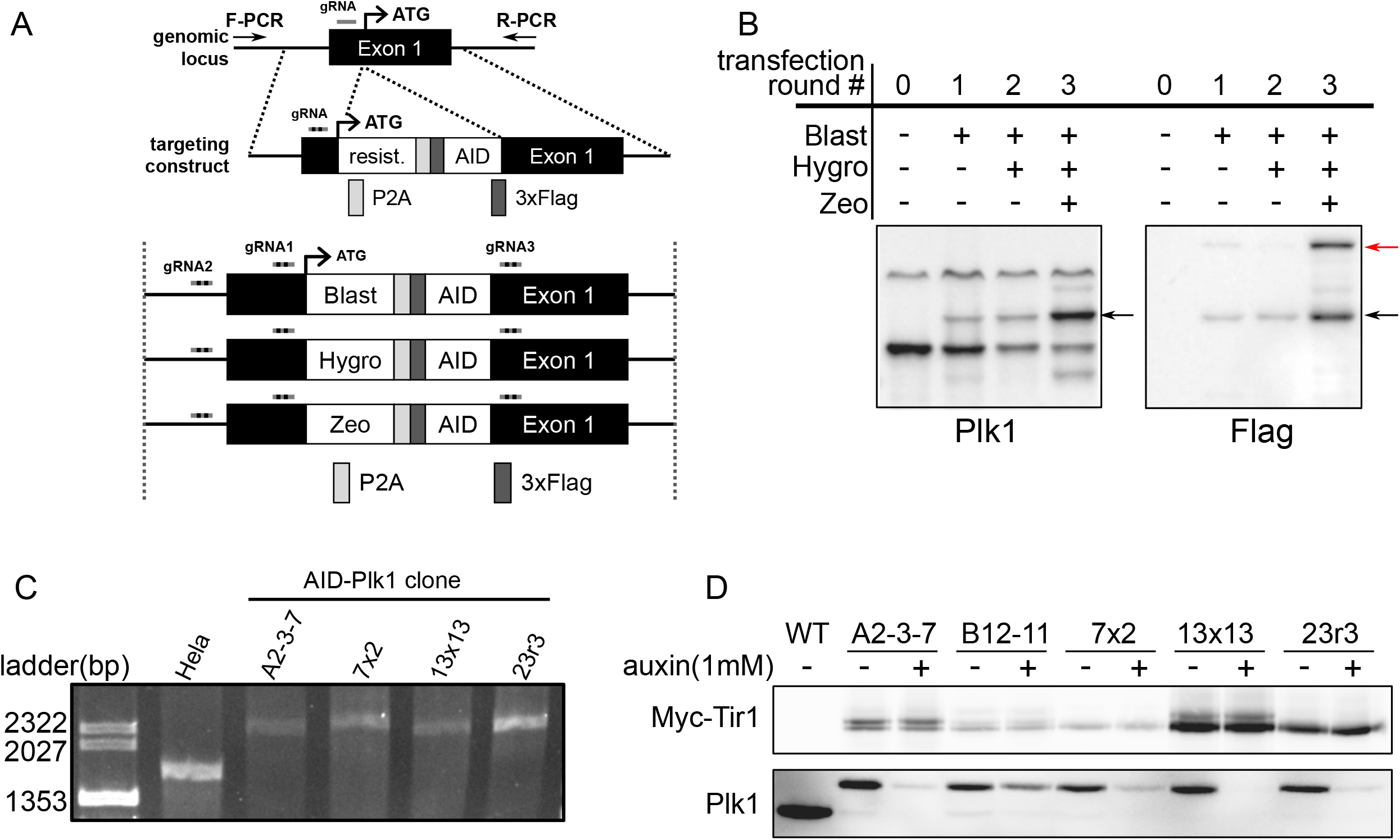
Generation of homozygous AID-Plk1 knock-in cells. (A) CRISPR-Cas9-based homologous recombination (HR) endogenous tagging scheme. gRNAs targeting the Plk1 locus adjacent to the start codon and HR templates containing an antibiotic resistance cassette, a P2A ribosomal skip site, a 3xFlag tag, and the AID tag for in-frame insertion to the N-terminus of Plk1. Blasticidin, hygromycin, and Zeocin resistance cassette-containing HR constructs include gRNA resistance mutations to three separate gRNAs, allowing for sequential transfection and selection with each HR construct/gRNA combination. (B) Pools of HEK293T CRISPR/Cas9 cells following sequential rounds of transfection and selection. Black arrows show the emergence of a greater proportion of AID-tagged Plk1 as transfection rounds progress by anti-Plk1 western blot. Red arrow shows Flag-tagged Cas9 by anti-Flag western blot. (C) Genomic PCR allows for confirmation of homozygous insertion of HR cassettes. (D) After Tir1 introduction, clones are screened by western blot for Tir1 expression and ability to degrade AID-Plk1 in the presence of auxin over 2 hours. Five representative AID-Plk1 degrading clones are shown.

### Individual AID-Plk1 knock-in HeLa clones exhibit variable rates of auxin-dependent AID-Plk1 degradation

We noted that among these clones (**Figure 3D**), the extent of degradation appeared highly variable within the 2-hour timeframe of initial screening. To quantitatively measure these differences, we performed an extended time course of auxin addition in slower (B12-11) and a faster (23r3) degrading clones and analyzed them by western blotting in biological triplicate (**Figure 4A**). Quantification of AID-Plk1 intensities over time revealed a Plk1 half-life in Taxol arrest of ∼ 31 minutes in the slower-degrading clone (**Figure 4B, blue line**). In contrast, the faster degrading clone exhibited an AID-Plk1 half-life of ∼ 8 minutes, a considerably faster auxin-dependent turnover rate that we observed much less frequently among AID-Plk1 knock-in clones that we screened (**Figure 4B, red line**). In addition, the overall extent of degradation was greater for the faster than for the slower clone (∼ 90% and 65% depleted at 3 hours, respectively). Auxin-dependent degradation rates under asynchronous conditions in which Plk1 expression is low were slightly lower than those observed in mitosis for both clones **(Figure S2)**.

**Figure 4.**
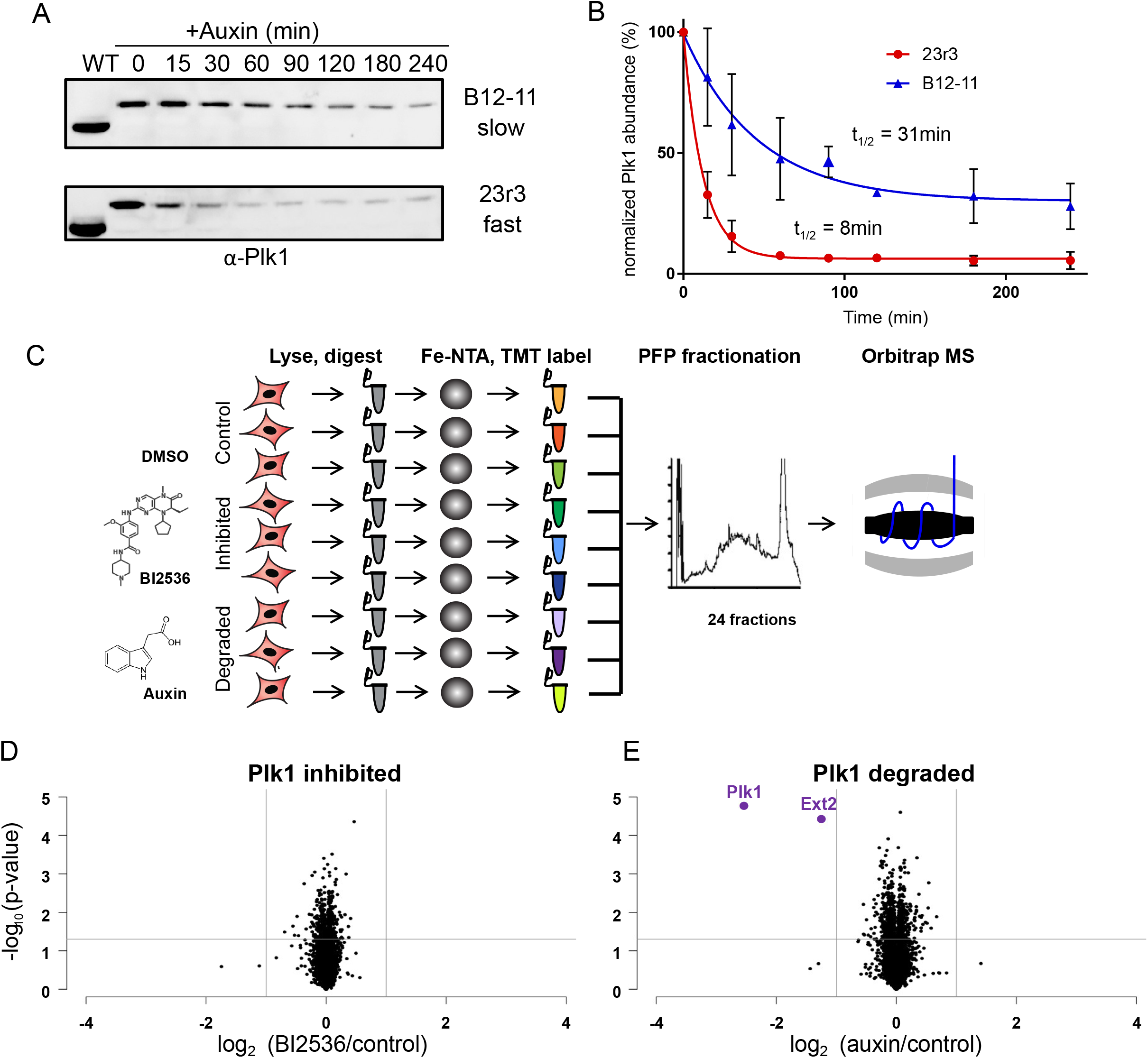
AID-Plk1 clonal degradation kinetics and TMT proteomics workflow. (A) Auxin time course of fast (23r3) and slow (B12-11) degrading AID-Plk1 clones by western blot. (B) Western blots from (A) in triplicate, quantified, and fit to an exponential decay for calculation of auxin induced AID-Plk1 half-lives. (C) Experimental scheme for 9-plex TMT proteomic and phosphoproteomic experiments of Taxol-arrested, AID-Plk1 clones: Triplicates of control treated cells, cells treated with Plk1 inhibitor BI2536 at 1μM for 1 hour, and cells treated with 1mM auxin for 2 hours. (D) Volcano plot of protein abundance changes in AID-Plk1 clone 23r3 upon BI2536 treatment (Plk1 inhibited). Each point represents one of 8349 proteins quantified. (E) Volcano plot of protein abundance changes in AID-Plk1 clone 23r3 upon auxin treatment; two proteins decreased significantly, Plk1 and Ext2.

### Auxin-dependent degradation via Tir1 is highly selective for AID-Plk1 protein in Taxol-arrested HeLa cells

We then sought to use these AID-tagged Plk1 cell lines in proteomic and phosphoproteomic experiments to determine the selectivity of auxin-dependent protein degradation in mitotic HeLa cells and to compare phosphorylation site abundance changes between acute cessation of Plk1 activity using BI2536, a highly selective Plk inhibitor, and auxin-induced loss of Plk1 protein. We designed a multiplex quantitative proteomics experiment in which the fast clone 23r3 was arrested in mitosis using Taxol and then either treated with auxin for 2 hours, BI2536 for 1 hour, or DMSO as a control (**Figure 4C**). These cells were treated in triplicate, lysed, and a portion of each lysate was analyzed for protein abundance by TMT labeling, peptide fractionation, and quantitative LC-MS/MS (**Figure 4C**). We identified and quantified 8349 proteins in this experiment, in which treatment of 23r3 cells with BI2536 showed no significant changes in protein abundance of 2-fold or greater when compared to control (**Figure 4D; Supplementary Table 2**). Treatment with auxin resulted in a 5.8-fold decrease in Plk1 abundance (**Figure 4E**), consistent with our results from western blotting (**Figure 4B**). Importantly, only one other protein, EXT2, a protein unrelated to Plk1 function in any known way, changed by more than 2-fold upon auxin addition. These data strongly support the assertion that the auxin-inducible degron tag, the core recognition elements of which (a WPPV sequence motif) do not occur in the human proteome, is a highly specific and selective substrate of the Tir1-containing human SCF E3 ubiquitin ligase complex in the presence of auxin in mitotic HeLa cells.

### Phosphoproteomic responses of faster degrading AID-Plk1 clones to auxin is highly correlated with responses to selective Plk1 inhibition with BI2536

We then processed the remaining 23r3 lysate digests for phosphoproteomics by phosphopeptide enrichment, TMT labeling, fractionation, and analysis by LC-MS/MS. We conducted an additional phosphoproteomics experiment with the slower degrading B12-11 clone to assess how protein turnover rate and extent of degradation might impact our ability to identify Plk1 candidate phosphorylation sites. In the analysis of the slower degrading B12-11 clones, we identified and quantified 23,072 phosphopeptides (**Supplementary Table 3**). Of these phosphopeptides, 422 were significantly reduced upon Plk1 inhibition with BI2536 (**Figure 5A**), while 246 were reduced upon auxin-induced degradation of AID-Plk1 (**Figure 5B**). However, in the analysis of the faster degrading 23r3 clone, from a total of 34,266 phosphopeptides quantified (**Supplementary Table 4**), a similar number were significantly reduced upon both Plk1 inhibition (486 phosphopeptides) (**Figure 5C**) and auxin-induced Plk1 degradation (472 phosphopeptides) (**Figure 5D**). Note that the difference in extent of phosphopeptide identification between these analyses is due to the use of Orbitrap Fusion and Orbitrap Fusion Lumos mass spectrometers. Intriguingly, despite the increased phosphoproteome coverage on the Orbitrap Fusion Lumos compared to the Orbitrap Fusion, a similar number of phosphopeptides were reduced upon the addition of the Plk inhibitor BI2536 in the slower B12-11 and faster 23r3 clone. However, the difference in response to auxin treatment in both cell lines suggests that the rate and degree of degradation impact the sensitivity of substrate identification. Motif analysis of significantly reduced phosphopeptides upon treatment with BI2536 or auxin in either fast or slow degrading cells revealed an enrichment of the canonical Plk1 substrate motifs [D/E/N]Xp[S/T][hydrophobic] and p[S/T]F (**Figure S3)**. Closer inspection of these auxin-responsive phosphoproteins identifies many previously identified Plk1 substrates, including 53BP1 at S1618 [45], Cdc27 at S426 and S435 [46], and Mklp2 at S528 [47], among others.

**Figure 5.**
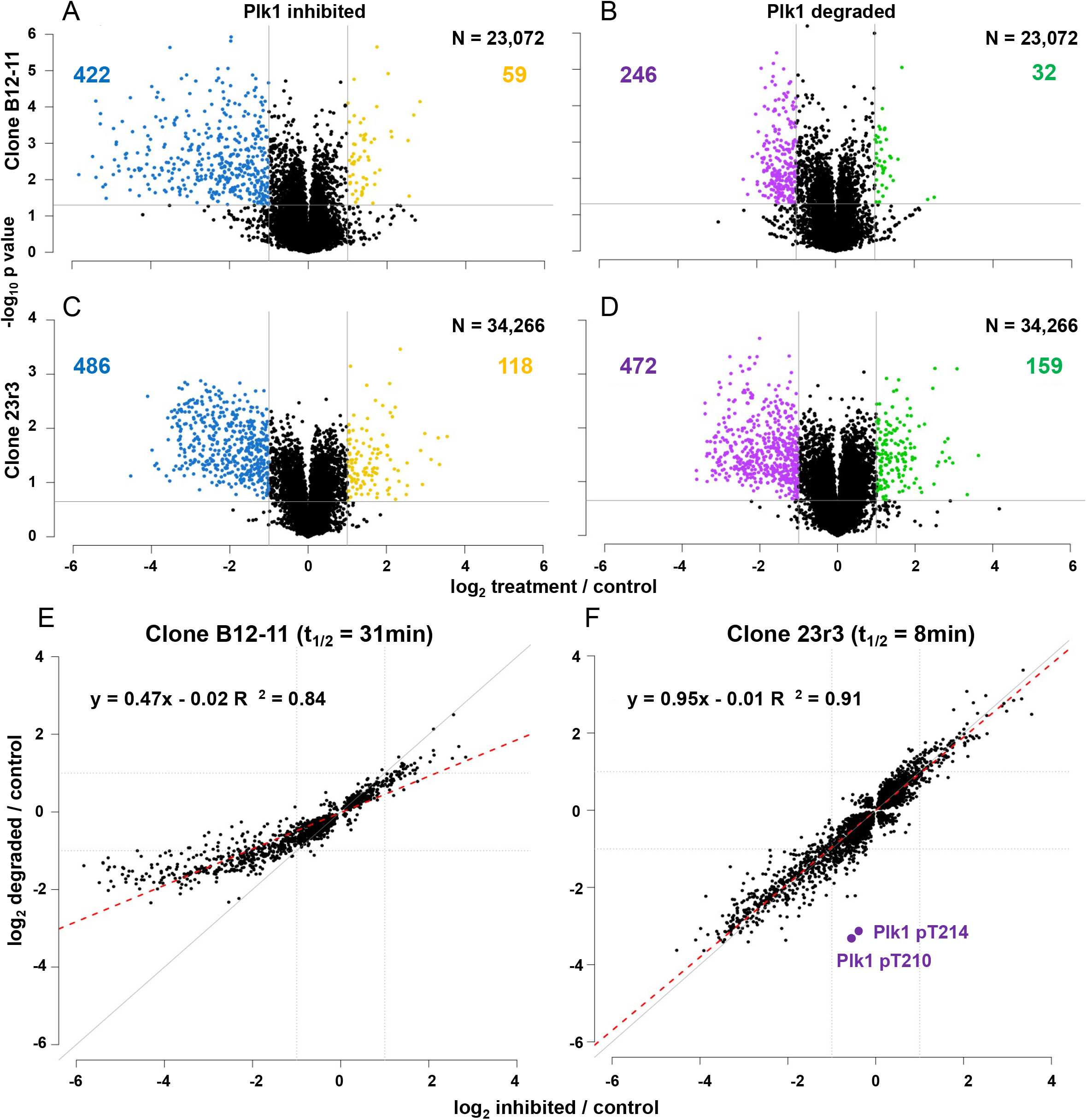
Quantitative phosphoproteomics of auxin-treated AID-Plk1 cells reveals candidate Plk1 substrates. Volcano plots of phosphosite abundance changes due to Plk1 inhibition (A) or AID-Plk1 degradation (B) in 23,072 phosphorylation sites from slow degrading B12-11 cells. Volcano plots of phosphosite abundance changes due to Plk1 inhibition (C) or AID-Plk1 degradation (D) in 34,266 sites from fast degrading 23r3 cells. Correlation plots of significant fold changes from both kinase inhibition vs degradation in B12-11 (E) and 23r3 (F) cells, including linear regression analysis (red dashed line). Auxin-dependent phosphosite changes from clone B12-11 correlate well with those due to inhibition but are decreased in magnitude. In contrast, auxin-dependent phosphosite changes from clone 23r3 are proportionally much closer to those due to inhibition. Note that two phosphosites that decrease exclusively in the degradation condition are sites on Plk1 itself.

To assess the degree to which both BI2536 and auxin downregulated individual candidate Plk1 substrate phosphopeptides, we extracted all phosphopeptides from either multiplex that exhibited a significant (p-value < 0.05) change in abundance in either BI2536- or auxin-treated cells and plotted these fold-changes against one another. While we also observed a high degree of correlation (R^2^ = 0.86) between both conditions in the slower degrading B12-11 clone, there was a compression of the magnitude of the fold-change difference in the auxin-treated cells (slope of correlation = 0.47) (**Figure 5E**). In the faster degrading 23r3 clones, however, we observed that phosphopeptide fold-changes upon auxin- or Plk inhibitor-treatment were both highly correlated (R^2^ = 0.91) and exhibited a magnitude of difference that is on average very consistent between them (slope of correlation = 0.95) (**Figure 5F**). Thus, treatment of Taxol-arrested HeLa cells with auxin does not appear to generate significant off-target signaling responses in these phosphoproteomes. In addition, while the fold-changes of most phosphopeptides were highly correlated between both conditions, two phosphorylation sites appear uniquely sensitive to auxin and not BI2536: Plk1 pT210 and pT214. Plk1 T210 is the activation T-loop phosphorylation site and is phosphorylated by Aurora kinase A [48, 49], while the kinase responsible for phosphorylating Plk1 T214 remains unknown. The non-correlated decreases in these phosphopeptides abundances are due to the degradation of the Plk1 protein upon auxin addition.

### Validation of novel Plk1 candidate substrates by *in vitro* kinase assays

While many of these candidate Plk1 substrate sites have been observed previously by us and by others, some of them are either new candidate sites on known Plk1 substrates or sites on proteins not previously linked to Plk1. To validate these novel sites, we cloned and purified four candidate substrate proteins involved in a diverse range of biological processes, which could indicate novel mitotic Plk1 activity: COIL, a component of nuclear coiled bodies thought to be regulated through mitotic phosphorylation [50-52]; DEK, a protein involved in chromatin organization that is also a potential oncogene [53, 54]; FAM54B, a mitochondrial fission regulating protein of unknown function; and NCK1, a cytoplasmic adaptor protein known to function in receptor tyrosine kinase signaling and the DNA damage response among others [55, 56]. These candidate substrates were expressed from 3xFlag expression constructs in HEK293T cells, purified, and used as substrates in *in vitro* kinase reactions with Plk1. To control for co-purifying kinases that may be in either the substrate or Plk1 preparations, we performed these reactions under four conditions: i) a negative control with the substrate of interest and no ATP (-ATP condition) to quantify basal phosphorylation of the purified proteins, ii) substrate protein and ATP (+ATP condition) to identify phosphorylation by co-purifying kinases in the isolated substrate preparation, iii) substrate protein, recombinant Plk1 and ATP, to identify candidate sites phosphorylated *in vitro* by Plk1 (+ATP+PLK1 condition), and iv) substrate protein, recombinant Plk1, ATP and BI2536 to identify co-purifying kinases in the recombinant Plk1 preparation (+ATP+PLK1+BI). *In vitro* kinase reactions were analyzed by label-free quantitative LC-MS/MS (**Figure 6A; Supplementary Table 5**). Candidate Plk1 substrates should have significantly higher phosphorylation site occupancy in the +ATP+Plk1 condition when compared to the other three control conditions. Several residues on COIL were identified as potential Plk1 substrate sites upon auxin addition by global phosphoproteomics: T289 or T290 and a cluster of sites spanning S571, S572, and S573. *In vitro* kinase reactions identified T290 and S571 with increased phosphorylation occupancy suggestive of Plk1 substrate sites (**Figure 6B**). S51 on the candidate substrate DEK was identified as auxin-responsive by phosphoproteomics and was also increased in occupancy in *in vitro* kinase reactions with purified Plk1 (**Figure 6C**). One site on NCK1, S96, was identified as a candidate Plk1 substrate site by phosphoproteomics; however, while its phosphorylation abundance in the +ATP+Plk1 condition was considerably higher than in control reactions lacking kinase, its abundance was partially increased in the +BI2536 inhibitor condition (**Figure 6D**). This suggests that NCK1 S96 might be a substrate of both Plk1 and a potentially co-purified kinase in the Plk1 preparation. Finally, the single candidate substrate site S261 on FAM54B was not observed in any *in vitro* kinase reaction condition **(Figure S4)**. In addition, the *in vitro* kinase reactions for all four candidate substrates yielded additional sites in the presence of Plk1 that were not identified in the global phosphoproteomics experiment (data not shown).

**Figure 6.**
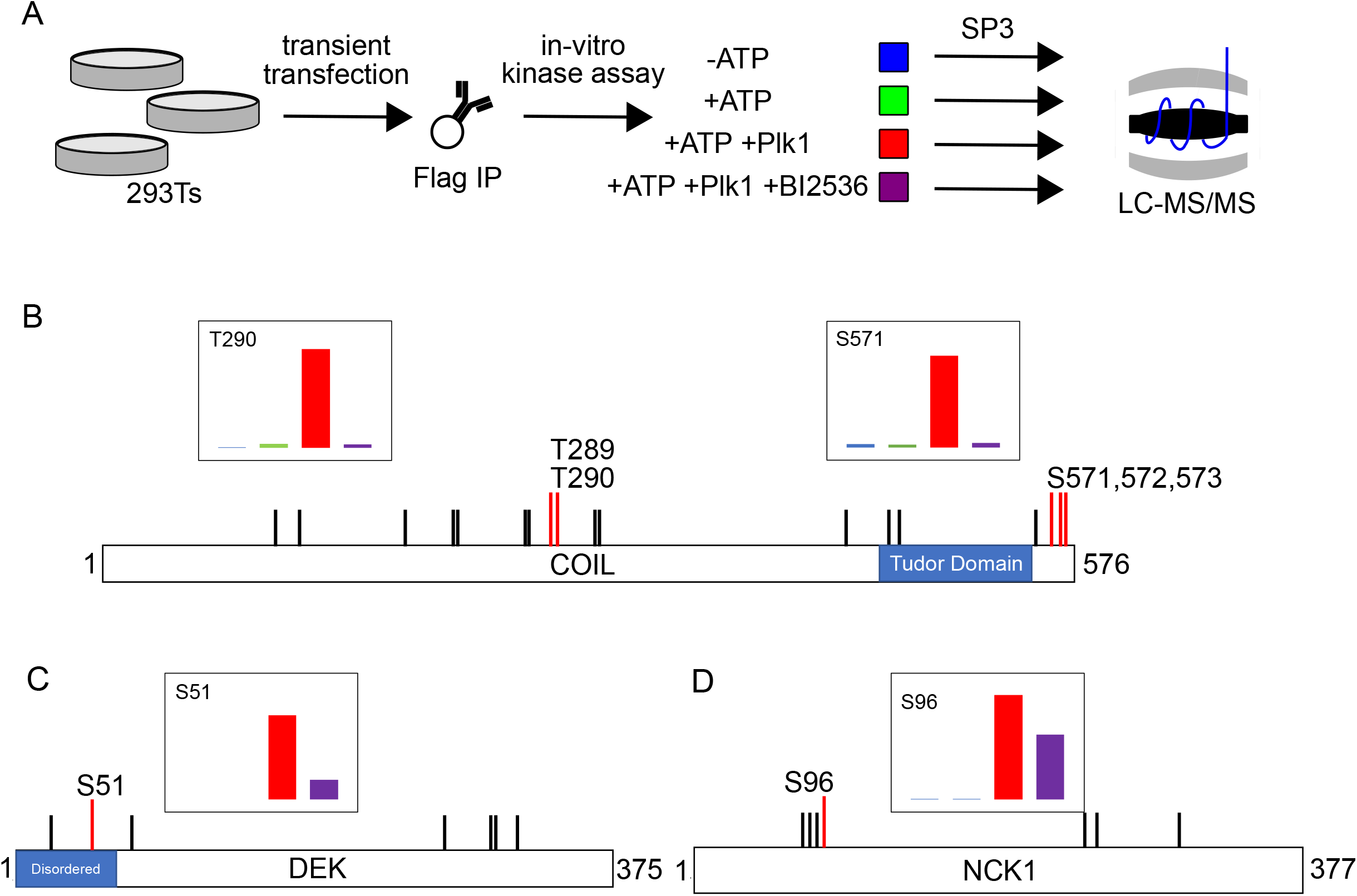
Candidate Plk1 substrate validation by *in-vitro* kinase assay. (A) Candidate Plk1 substrate validation scheme. 3xFlag tagged proteins were purified from transient expression in 293s. Proteins were incubated: alone (blue); with ATP (green); with ATP and purified Plk1 (red); or with ATP, Plk1, and BI2536 (purple). Proteins were then digested to peptides and analyzed by label-free quantification LC-MS/MS. (B) Schematic depiction of protein COIL. Lines show coverage of phosphorylation sites from LC-MS/MS analysis. Red lines represent candidate Plk1 substrate sites, black lines represent non-Plk1 sites. Inset bar graphs represent quantification of phosphorylation sites identified in the 4 conditions of the *in-vitro* kinase assay. Colors correspond to conditions in (A); T290 and S571 results corroborate results from AID-Plk1 phosphoproteomics. (C) Protein DEK depicted as in (B); *in vitro* kinase assay results for S51 are consistent with AID-Plk1 phosphoproteomics. (D) Protein NCK1 depicted as in (b); *in vitro* kinase assay result for S96 is supportive but not conclusive of it as a novel Plk1 substrate site.

### Proteome-wide steady-state protein turnover rates are modestly higher in faster versus slower degrading AID-Plk1 clones

Our analysis of auxin-dependent changes in the phosphoproteomes of faster degrading (23r3) and slower degrading (B12-11) HeLa cell lines revealed a strong dependence on this turnover rate and the degree of degradation. While the faster degrading cell line represents an ideal comparison to chemical inhibition, we found that such cell lines are difficult to generate. We reasoned that identifying the biological basis for the differences in turnover rate may reveal additional strategies to generate AID-kinase cell lines with inherently faster auxin-dependent degradation rates. For example, it could be possible that differences in turnover rate could be proportional to the abundance of Tir1 in individual clones. While Tir1 abundance appears to contribute to AID-Plk1 turnover rates, we found that clones with similar levels of Tir1 can have different AID-Plk1 degradation rates upon auxin addition.

Alternatively, we reasoned that differences in global protein turnover rates could be the underlying basis for the slower and faster AID-Plk1 degradation rates. To test this, we treated B12-11 and 23r3 cells in biological triplicate with cycloheximide (CHX) to halt new protein synthesis, collected cells at four-hour and 20-hour timepoints after CHX addition and compared their protein levels to untreated control cells by quantitative proteomics (**Figure 7A; Supplementary Table 6**). Estimates of protein turnover rates were calculated by taking the proportion of protein abundance that decreased between timepoints divided by the time between timepoints and averaged across the three replicates. The distribution of turnover rates demonstrates that proteome-wide protein turnover in 23r3 cells is faster than in B12-11 cells and is especially prominent in the first four hours post-CHX addition (**Figure 7B**). To ascertain if this difference between cell lines is consistent on a per-protein basis, we compared normalized protein abundances for each protein from B12-11 and 23r3 cells at the four (**Figure 7C**) and 20-hour timepoints (**Figure 7D**). Individual protein turnover rates for most proteins appear higher in 23r3 cells, with the degree of turnover rate difference being greater at the four-hour than at the 20-hour timepoint. Although beyond the scope of this experiment, we speculate that this compression of protein abundances at 20 hours may reflect cellular responses to prolonged CHX, rather than ongoing dynamic protein turnover. However, we note that while global protein turnover is indeed faster in 23r3 cells, this difference in abundance is on average roughly 21% at four hours. These data suggest that the auxin-induced degradation differences between the faster 23r3 and the slower B12-11 clones may in part be explained by differences in total protein turnover machinery between them.

**Figure 7.**
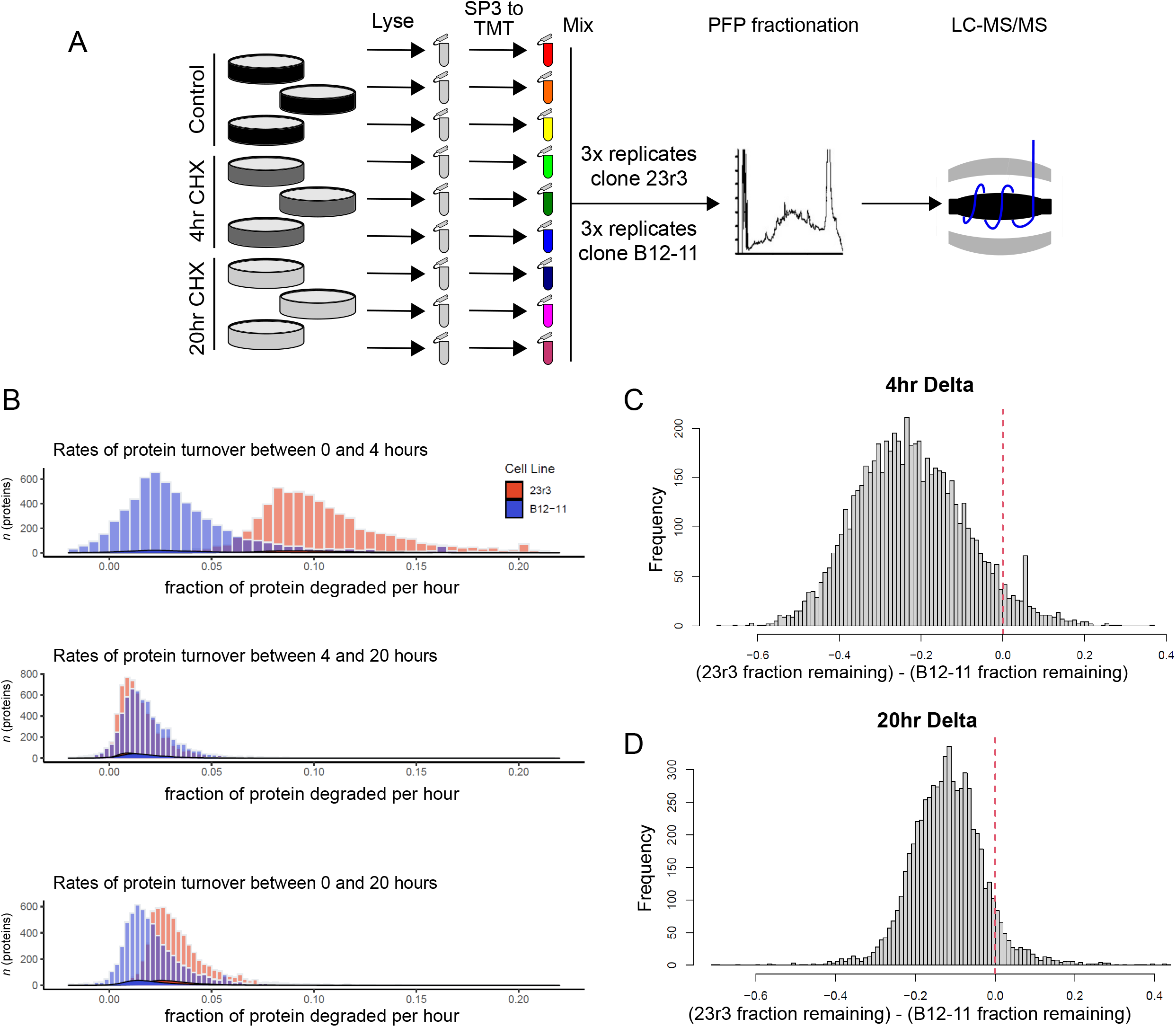
Quantitative proteomics of cycloheximide-treated cells reveals basal protein turnover rate differences between clones. (A) TMT 9-plex proteomics workflow for triplicates of AID-Plk1 fast (23r3) and slow (B12-11) degrading cells treated with vehicle and cycloheximide for 4 and 20 hours. (B) Distribution of rates of normalized protein abundance turnover per hour between timepoints 0 and 4 hours (top), 4 and 20 hours (middle), and 0 and 20 hours (bottom). (C, D) Distributions of per protein abundance differences (deltas) in clone 23r3 minus clone B12-11 at 4 hours (C) and 20 hours (D) after cycloheximide addition. Negative delta values indicate proteins with greater fraction of protein remaining in clone B12-11 than in clone 23r3 at a specific time after cycloheximide addition.

## Discussion

Linking protein kinases to their direct protein substrates provides candidate mechanisms of function that relate kinase activity to distinct molecular signaling pathways that can result in activation, inhibition, localization, aggregation, dissociation, degradation, or other aspects of substrate protein function. While modern technologies and approaches in quantitative proteomics have enabled the routine identification of many thousands of phosphorylation sites, providing biological context for these sites remains challenging. In this work, we sought to provide a generalized roadmap for determining kinase-substrate relationships that could pave the way for future studies to illuminate the “dark” kinome. We chose Plk1 as a model system for developing this technology, given its well-characterized roles in cell division and cancer, and the availability of a highly selective inhibitor for validation purposes. Inducible protein degradation technology continues to evolve and develop and is being adapted to solve many problems in biology. We have shown in this work that AID-tagged kinases can be used in conjunction with quantitative proteomics and phosphoproteomics to uncover candidate substrates. As part of our rigorous validation, we demonstrated that shorter AID tags did not interfere with the kinase activity, substrate preference, interactions, or localization of Plk1. While conceptually straightforward, the use of CRISPR-Cas9 to create homozygous knock-in clones presented additional technical challenges that required a sequential knock-in and selection system to enable the progressive enrichment of tagged proteins in pools of cells, with the need to isolate and screen individual clones once to find homozygotes. The process remains time intensive but advances in screening for clones by single cell sorting with fluorescent protein reporters may provide avenues to speed up this process.

During our testing of different homozygous AID-Plk1 cell lines, we observed a wide range in the rate and overall extent of auxin-induced degradation, which could not be predicted entirely by Tir1 abundance levels alone. In comparing cell lines with slow and fast Plk1 degradation kinetics, we noted that while the slower degrading cell line treated with auxin induced the same general biological response as chemical inhibition, the magnitude of this response was suppressed compared to chemical inhibition and the faster degrading cell line. This might be due to differences in the mechanism between loss of kinase activity in a chemical inhibition system and a degradation system. Upon inhibitor addition to cells, the entire complement of kinase molecules is rapidly inhibited in proportion to the intracellular concentration of the inhibitor and its target affinity. In contrast, initiating the degradation of a kinase will result in protein turnover at a rate consistent with its abundance in the cell. This means that during the course of degradation, kinase activity decreases but persists from the pool of kinase that has yet to be degraded. Thus, fast degradation kinetics are essential to evoke phosphoproteome responses comparable to kinase inhibition. This suggests that loss of kinase protein by degradation to a minimal threshold, followed by a period of lack of kinase activity, is required for phosphatase-dependent turnover of their substrate sites to reveal them as such by quantitative phosphoproteomics. Additionally, it is important to emphasize that while faster degrading cell lines do function better at substrate discovery, and that efforts should be made to identify and use faster degrading clones, even slower degrading cell lines can still uncover real kinase-specific cell biology, albeit at the cost of depth of coverage and false negative identifications. This should inspire confidence towards adapting this technology towards all understudied kinases that lack specific chemical inhibitors, where those tools are not yet available to validate substrates discovered from AID-kinase screens. Finally, we note that while our correlation analysis between Plk1 inhibitor and AID-Plk1 degradation was critical in validating AID-Plk1 as a viable strategy for identifying kinase-substrate relationships, future drug discovery efforts may find value in validating novel chemical inhibitors as selective for their target of interest through phosphoproteomic comparison with AID-target kinase cell lines.

In an effort to gain insights into the reasons behind the difference in induced degradation kinetics between fast and slow degrading clones, we assessed differences in basal protein turnover rates between two of them. Interestingly, there does appear to be an increased rate of turnover of most proteins in the faster AID clone when compared to the slower clone, albeit a modest one. This observed difference in basal protein turnover rate in our estimation likely does not fully explain the difference in induced protein degradation rates, so it will also be interesting to explore additional factors that may affect Tir1-specific function. Until it is possible to modulate these cell-intrinsic factors to routinely create fast degrading cell lines, more work is required to develop strategies for screening out degrading cell lines.

While the goal of this work was to establish auxin-inducible degradation technology for phosphoproteomics to uncover substrates of kinases, we were also able to conduct deep phosphoproteomics experiments of phosphosite abundance changes due to loss of Plk1 specifically. This has generated over 500 candidate substrate sites for Plk1 in mitosis. Many of these sites are not observed in prior screens for Plk1 substrates. These new sites represent a wide range of biological functions that expands the roles that Plk1 plays in facilitating diverse aspects of cell division. Candidate sites on proteins such as COIL and FAM54B suggest a role for Plk1 in triggering the fission of organelles and other large macromolecular structures at the onset of mitosis. Confirming DEK as a Plk1 substrate may provide further mechanistic insights into how Plk1 functions in a cancer context, as DEK is an emerging oncogene in a multitude of cancers and an increase in Plk1 activity has been observed in many cancers as well. In addition to the phosphorylation sites we validated here, other significant substrates from the screen may provide new mechanistic insights into Plk1’s role in the DNA damage response as well as fine-tune its more established roles in spindle assembly and organization and in centrosome regulation.

In conclusion, we believe this work lays the foundation for adapting inducible degradation technology to shed light on the mechanisms that underlie kinase functions that have gone understudied due to a lack of appropriate tools. The specificity, selectivity, and kinetics of inducible degradation allow for an accurate assessment of kinase substrates and are expected to enable new insights into the complex biology of the phosphoproteome.

## Acknowledgements

The authors would like to thank members of the Gerber and Kettenbach labs for helpful discussions. We specifically would like to thank Andrew Holland for helpful discussions and insights into auxin inducible degradation. This work was supported financially by grants from the National Institutes of Health (NIH) R01GM122846 and S10OD016212 to S.A.G., R35GM119455 to A.N.K, and shared resource support from P30CA023108 to the Norris Cotton Cancer Center.

## Supporting Information

**Supplementary Table 1:** LC-MS/MS data from replicate HA-Plk1 and HA-AID-Plk1 immunoprecipitation.

**Supplementary Table 2:** TMT proteomics data for protein abundances in fast degrading 23r3 cells treated with vehicle, Plk inhibitor BI2536 and auxin.

**Supplementary Table 3:** TMT proteomics data for phosphopeptide abundances in slow degrading B12-11 cells treated with vehicle, Plk inhibitor BI2536 and auxin.

**Supplementary Table 4:** TMT proteomics data for phosphopeptide abundances in fast degrading 23r3 cells treated with vehicle, Plk inhibitor BI2536 and auxin.

**Supplementary Table 5:** LC-MS/MS data from *in vitro* kinase assays with purified Plk1 and candidate substrates COIL, DEK, FAM54B and NCK1.

**Supplementary Table 6:** TMT proteomics data for protein abundances in fast degrading 23r3 and slow degrading B12-11 cells treated with vehicle or cycloheximide for 4 and for 20 hours.

---

**Figure S1:**
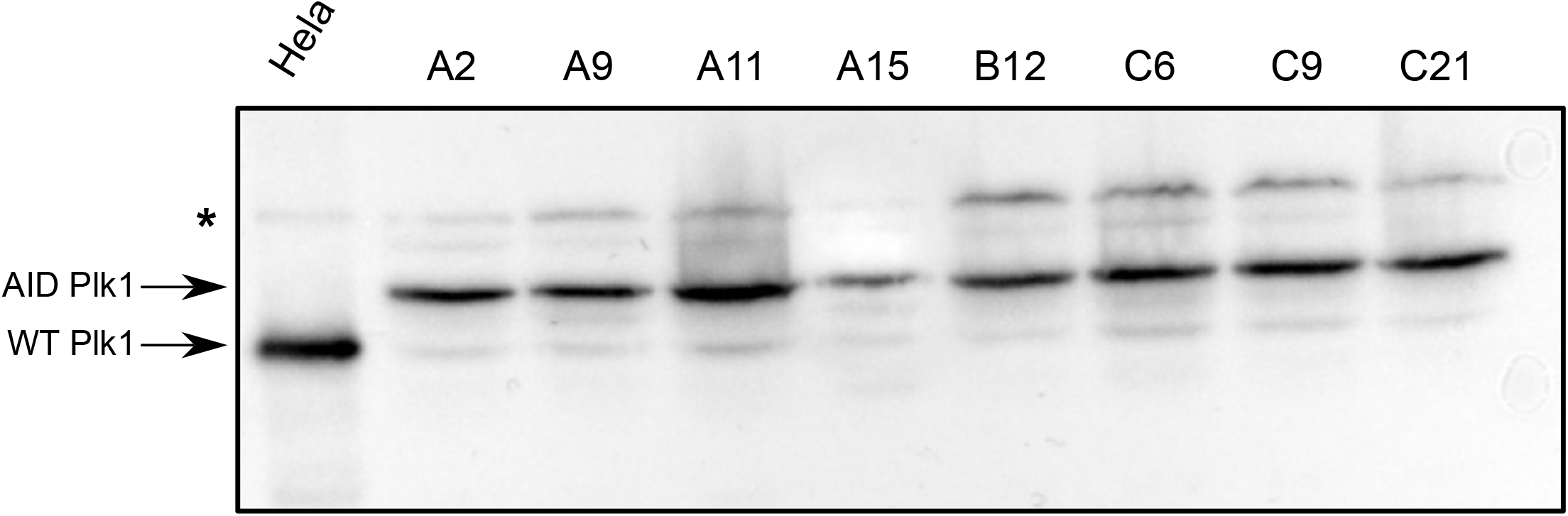
Western blot of representative AID-Plk1 clones. Plk1 western blot of pre-screened clones demonstrating homozygosity of AID-Plk1 insertion. Arrows indicate relative mobility of wild-type Plk1 and AID-Plk1. Asterisk denotes non-specific band from anti-Plk1 antibody.

**Figure S2:**
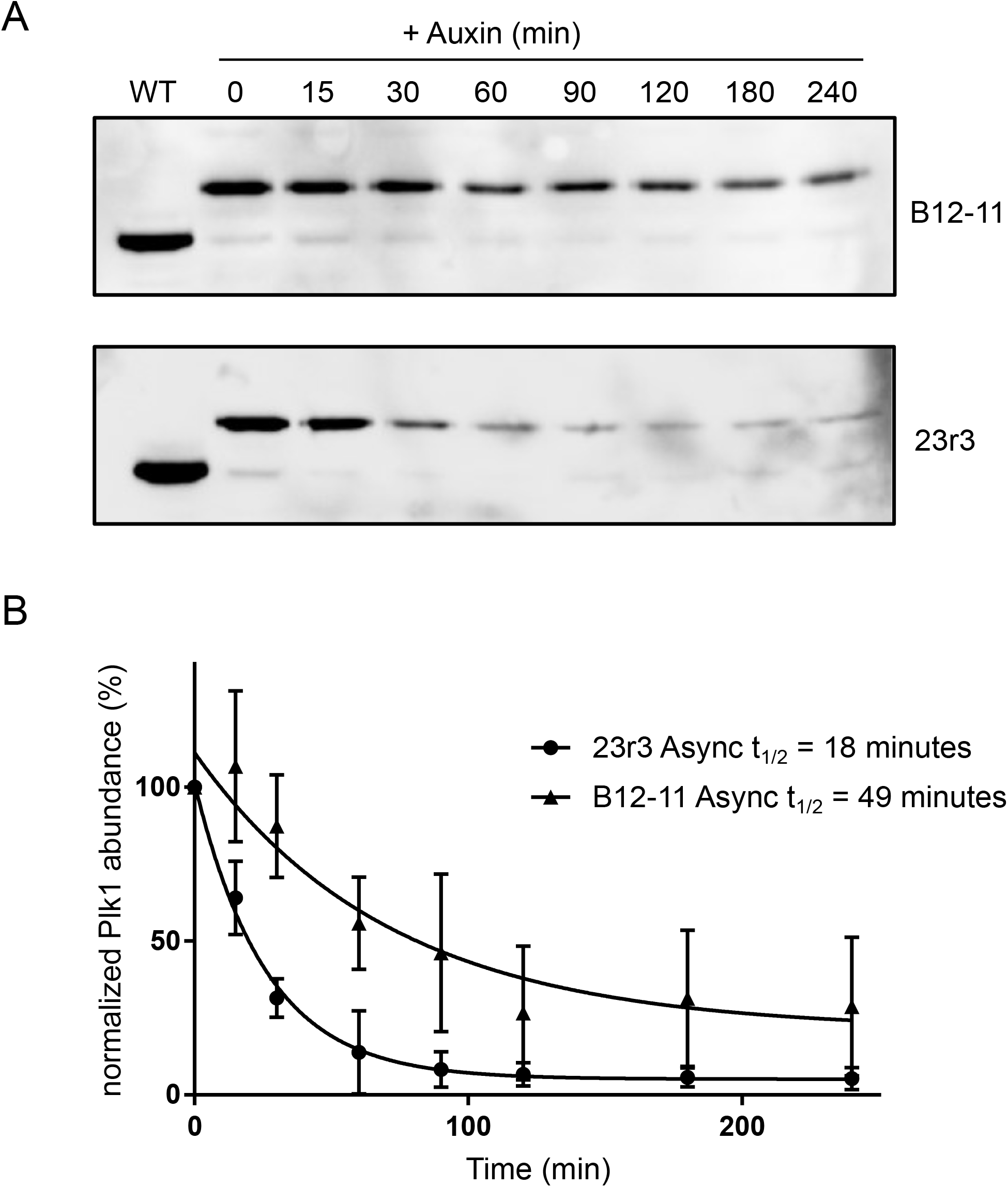
Asynchronous AID-Plk1 degradation kinetics. (A) Representative western blots of a time course for clone B12-11 and 23r3 after 1mM auxin addition in asynchronous cells. Time course and western blots were repeated in triplicate. (B) Quantification of western blots from (A). Protein half-lives were calculated by fitting exponential regressions to averages from triplicate quantifications. Error bars show sample error.

**Figure S3:**
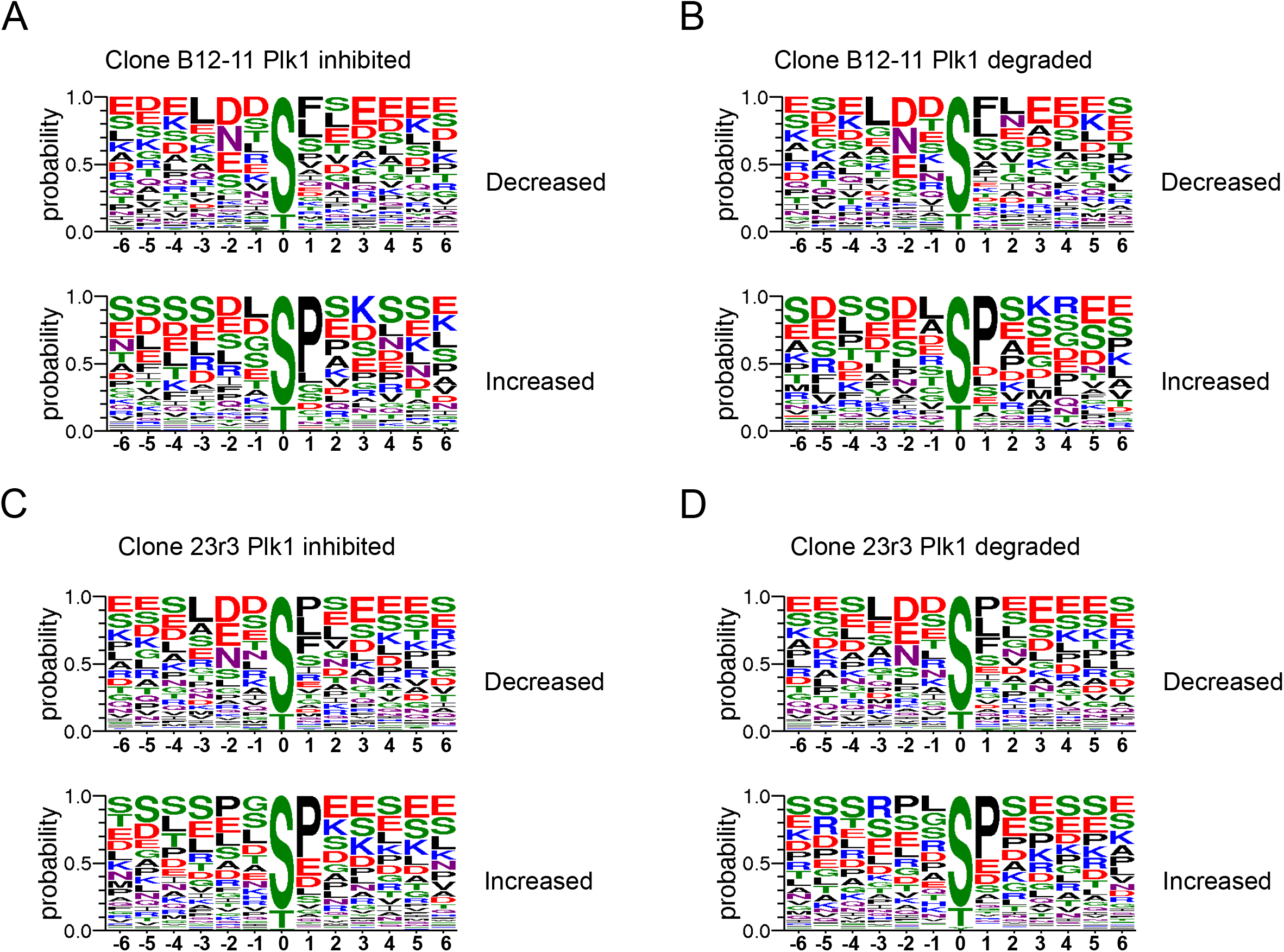
Phosphorylation motifs from AID-Plk1 phosphoproteomics experiments. Weblogo representation of phosphorylation motifs generated from significantly significant decreasing (top) and increasing (bottom) sites that changed two-fold or more in (A) clone B12-11 with one hour BI2536 treatment, (B) clone B12-11 with two hour auxin treatment, (C) clone 23r3 with one hour BI2536 treatment, and (D) clone 23r3 with two hour auxin treatment.

**Figure S4:**
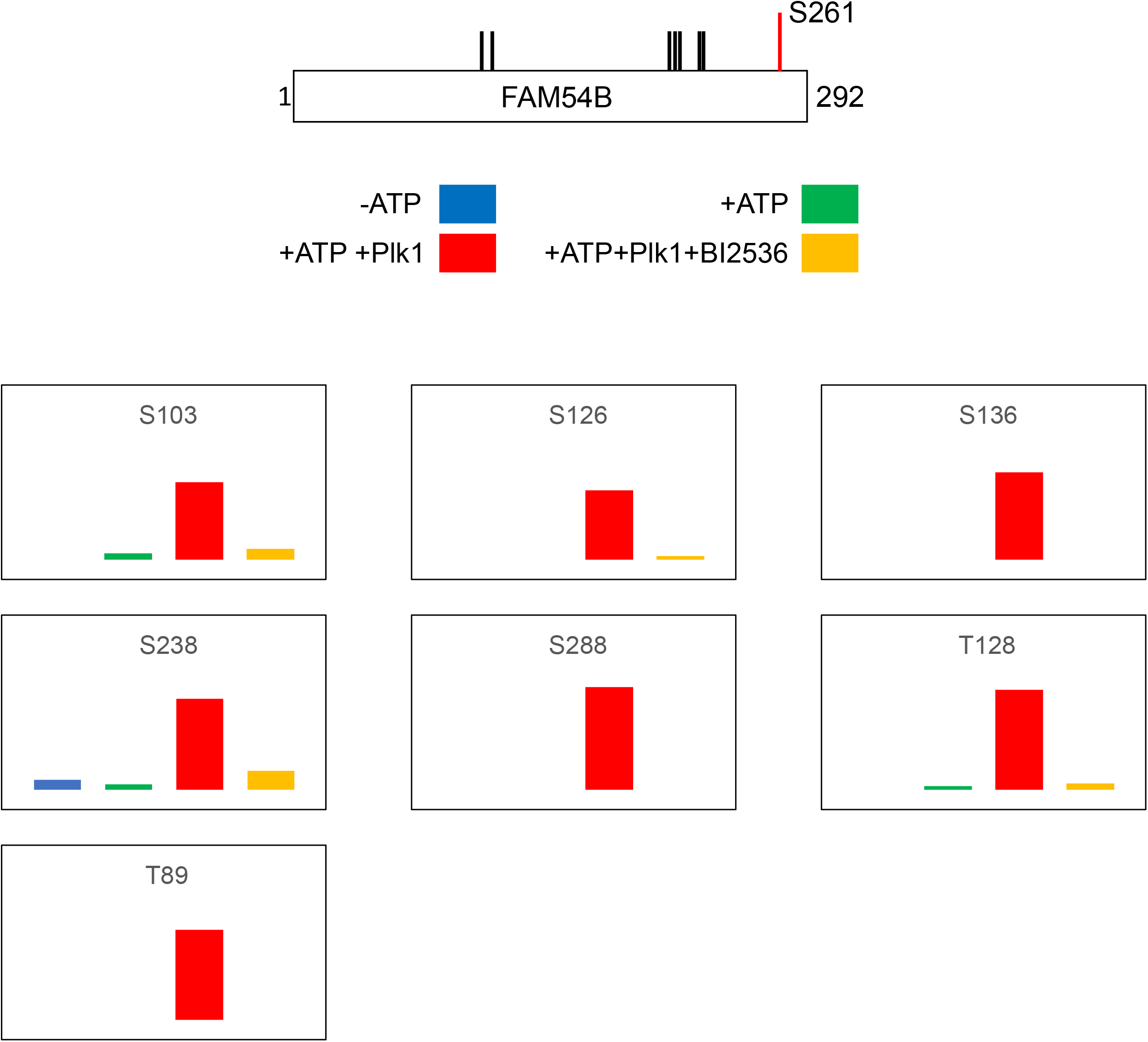
*In vitro* kinase assay of FAM54B with Plk1. All quantified phosphorylation sites detected from FAM54B *in vitro* with affinity-isolated FAM54B and recombinant Plk1. (top) Cartoon depiction of FAM45B with phosphorylation site coverage from phosphoproteomics experiment depicted with vertical lines. S261 is marked in red as the candidate Plk1 substrate site. (bottom) Graphs of LC-MS/MS quantifications of the respective phosphorylation sites detected in the Plk1 *in vitro* kinase assay. Colors represent conditions as per the key above. Lack of column represents missing quantification in that condition.

